# Wide field-of-view volumetric imaging by a mesoscopic scanning oblique plane microscopy with switchable objective lens

**DOI:** 10.1101/2020.06.29.177782

**Authors:** Wenjun Shao, Kivilcim Kilic, Wenqing Yin, Gregory Wirak, Xiaodan qin, Hui Feng, David Boas, Christopher V. Gabel, Ji Yi

**Affiliations:** Department of Medicine, Boston University School of Medicine, Boston Medical Center, Boston, USA; Neurophotonics Center, Boston University, Boston, USA; Renal Section, Department of Medicine, Boston University School of Medicine, Boston, USA; Department of Physiology and Biophysics, Boston University, Boston, USA; Departments of Pharmacology & Experimental Therapeutics and Medicine, Boston University, Boston, USA; Department of Biomedical Engineering, Boston University, Boston, USA; Department of Electric and Computer Engineering, Boston University, Boston, USA

**Author notes:** **Correspondence:** Ji Yi, Boston University, 650 Albany Street, 02118, Boston, USA. Department of Biomedical Engineering and ophthalmology, Johns Hopkins Univeristy, Baltimore, USA.

**Keywords:** Mesoscopic scanning oblique plane microscopy, wide field-of-view, volumetric imaging, fluorescence imaging

## Abstract

Conventional light sheet fluorescence microscopy (LSFM), or selective plane illumination microscopy (SPIM), enables high resolution 3D imaging over a large volume by using two orthogonally aligned objective lenses to decouple excitation and emission. The recent development of oblique plane microscopy (OPM) simplifies LSFM design with only one single objective lens, by using off-axis excitation and remote focusing. However, most reports on OPM has a limited microscopic field of view (FOV), typically within 1×1 mm^2^. Our goal is to overcome the limitation with a new variant of OPM to achieve mesoscopic FOV. We implemented an optical design of mesoscopic scanning OPM to allow using low numerical aperture (NA) objective lens. The angle of the intermediate image before the remote focusing system was increased by a demagnification under Scheimpflug condition such that the light collecting efficiency in the remote focusing system was significantly improved. We characterized the 3D resolutions and FOV by imaging fluorescence microspheres, and demonstrated the volumetric imaging on intact whole zebrafish larvae, mouse cortex, and multiple *Caenorhabditis elegans (C*. *elegans*). We demonstrate a mesoscopic FOV up to ~6× 5×0.6 mm^3^ volumetric imaging, the largest reported FOV by OPM so far. The angle of the intermediate image plane is independent of the magnification. As a result, the system is highly versatile, allowing simple switching between different objective lenses with low (10x, NA 0.3) and median NA (20x, NA 0.5). Detailed microvasculature in zebrafish larvae, mouse cortex, and neurons in *C. elegans* are clearly visualized in 3D. The proposed mesoscopic scanning OPM allows using low NA objective such that centimeter-level FOV volumetric imaging can be achieved. With the extended FOV, simple sample mounting protocol, and the versatility of changeable FOVs/resolutions, our system will be ready for the varieties of applications requiring *in vivo* volumetric imaging over large length scales.

## 1 Introduction

Light sheet fluorescence microscopy (LSFM), or selective plane illumination microscopy (SPIM) has been an essential imaging modality in life science (1–3), enabling high resolution 3D imaging over a large volume. The typical configuration of LSFM/SPIM has two orthogonally aligned objective lenses to decouple the excitation and collection by separate optical paths. As the illuminated light sheet section is entirely collected with high efficiency, LSFM ensures low photodamage and low photo-toxicity. However, due to the orthogonal arrangement of two objective lenses, the imaging space for the sample is limited, which makes it difficult for large samples and being integrated with conventional microscopic platforms. In addition, conventional LSFM compiles volumetric dataset by mechanically translating the inertia components, such as an objective lens or samples, which can be challenging for high-speed imaging on large volumes. Inverted and ‘Open-top’ LSFM/SPIM improve the sample mounting protocols by rotating both of the objective lenses to the opposite side of the sample but still need mechanical translation (4–6). The adoption of the focus tunable lens and piezoelectric actuator makes the z-stacking process faster, yet still being limited within 1Hz volume rate when imaging zebrafish brain (7,8).

To improve the resolution, increase volumetric imaging speed and enable flexible sample mounting/positioning protocols, different variants of LSFM /SPIM have emerged over the recent two decades (9–15). Among them, oblique plane microscopy (OPM) provides an attractive optical design and provides excellent balance among the above three aspects. In contrast to conventional LSFM that has two objective lens, OPM uses only single objective lens but applies an off-axis oblique light sheet excitation and a remote focusing system to capture the light sheet in a near-orthogonal angle (9–11,13,14). This “*single objective lens layout*” of an OPM liberates the sample placing space, and can be set up in the existing inverted, or upright microscopes. The first embodiment of OPM is developed by Dunsby *et al.* with mechanical translation of the sample. Swept confocally-aligned planar excitation (SCAPE) microscopy innovatively introduced scanning and descanning imaging strategy (16), enables completely translationless three-dimensional imaging. SCAPE demonstrated high-speed volumetric imaging of up to 200 volume per second (VPS) in the intact living samples (17). To increase the light collecting efficiency and improve the resolution, water immersion technique can be introduced in the remote imaging system (18). A tilt-invariant scanned OPM (SoPi) was investigated (19); two-photon excitation technique was adopted (20,21); and simultaneous multimodal 3D imaging with optical coherence tomography (OCT) was demonstrated (22).

So far, the existing OPM studies largely relied on a high numerical aperture (NA) objective lens, which limits the achievable FOV to a microscopic level typically <1×1 mm^2^. A diffractive OPM circumvented this constraint by redirecting the oblique image plane with a diffraction grating. As a result, it could achieve a larger FOV with the help of a low NA objective lens (23). Nonetheless, it still has a limit for the light collection because the diffraction grating has to be placed in the restricted space between two lenses in the intermediate image space. Here, we leverage a seemingly counterintuitive phenomenon that the angle between the excitation and detection light paths does not need to be near-orthogonal (22), and the tilted angle for the intermediate image plane is maintained with varying magnifications, simply under the Scheimpflug condition (24). As a result, our OPM setup is highly versatile in using a low NA objective lens to obtain mesoscopic FOV up to ~5×6 mm^2^, the largest one reported by OPM so far. We demonstrated using 10X 0.3NA and 10x 0.5NA objective lens to imaging different biological specimens crossing a large range of length scales, from sub-millimeter *C. elegans*, whole zebrafish larvae, to a mouse cortex in centimeter scale.

## 2 Materials and Methods

### 2.1 Experiment setup

Our design derived from our previous work on oblique scanning laser ophthalmoscopy (oSLO) that essentially implements scanning OPM using the natural ocular optics (25). We previously simulated the diffraction-limited 3D resolutions at varying NA of the objective lens from 0.1 to 0.9 (22), and demonstrated oSLO in rodent *in vivo* and in human *in vitro* (25–27). Here we implemented and remodel the optical design used in oSLO in a microscopic setting. The schematic layout of the experimental setup is shown in Fig. 1(a). The objective lens OL1 is a low NA objective lens, which makes it possible for a wide FOV in OPM.

**Fig. 1.**
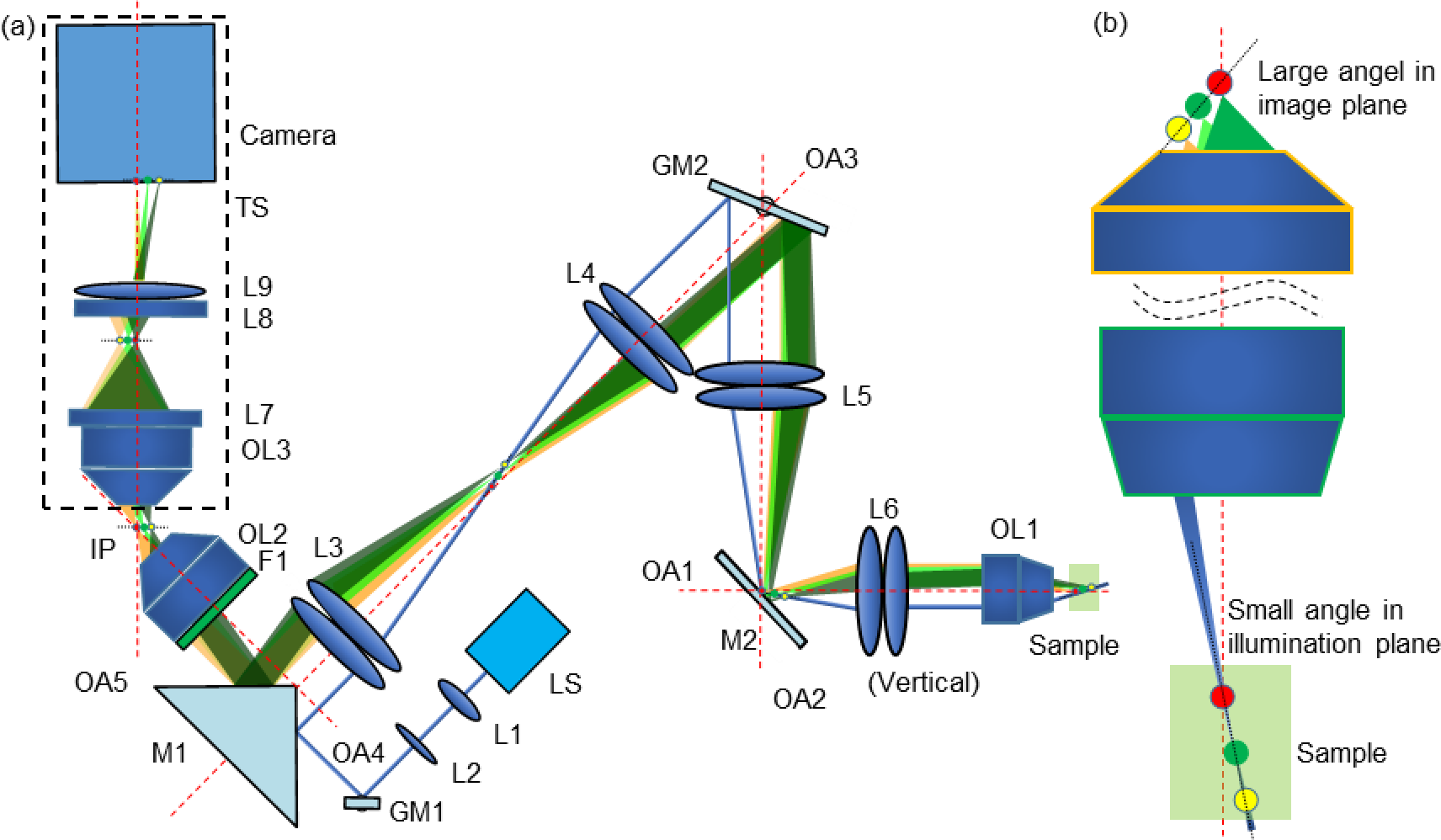
The experiment setup of mesoscopic OPM. a) The system schematic. L: lens; OL: objective lens; F: filter; IP: intermediate image plane; M: mirror; GM: galvanometer mirror; LS: light source, OA: optical axis, TS: translation stage. b) The angle enlargement of the image plane.

As shown in Fig. 1(a), excitation light is marked by blue, and emission light is marked by green and yellow. The excitation light was generated by a 488nm laser (LS), first coupled into single-mode fiber and then collimated by lens L1 (*f* = 10 mm). An additional focus-tunable lens L2 (Edmund: EL-3-10-VIS-26D) was used to make fine adjustments such that excitation and emission light could be confocal over a large FOV. The collimated light was first scanned by GM1 (Thorlabs: GVS201) and then redirected into a 1:1 relay lens group (L3-L4) by a right-angle prism mirror M1 (MRAK25-P01). As GM1 was placed at the focal plane of L3, a stationary scanning point was generated at the pupil plane of objective lens OL1 after two relay lens groups (L3: f = 100 mm, L4: f = 100 mm; L5: f = 100 mm, L6: f = 50 mm), and then a scanned light-sheet was created within the specimen. To maximize the angle of the oblique light-sheet, the excitation light was offset from the optical axis OA1-OA3 and incident on the edge of OL1. Another galvanometer GM2 (Nutfield: QS-12 OPD, 20 mm aperture), which was conjugated to the pupil plane of the OL1, was used to sweep the oblique light-sheet through the sample. As GM2 steered the excitation light-sheet and created a moving light-sheet within the sample, fluorescence emission mapped back on the same GM2 could be descaned. As a result, an intermediate stationary image plane (IP) could be created between OL2 (UPLSAPO 20×/0.75) and OL3 (UplanFL20×/0.5). The angle between the optical axis OA2 and OA3 was minimized to facilitate a more effective descanning over a large FOV. M1 was used to direct the fluorescence into OL2. An emission filter F1 (500-550nm) was placed in the back of OL2. The magnification of these two relay lens groups (L3-L6) was chosen to maximize the use of the numerical aperture of OL2. The lateral magnification *M*_*lateral*_ from the sample plane to the conjugated IP is designed to be less than 1. As the angle of the IP was significantly increased by the demagnification design, the remote imaging system could achieve sufficient light collecting efficiency. A translation stage TS of 4 degrees of freedom (X, Y, Z, and Yaw) was used to adjust the position and the angle of the remote imaging system.

OL1 can be switched between different low NA objective lenses, such as UplanFL10×/0.3 and UplanFL20×/0.5 from Olympus, to change FOVs and resolutions. The focal length of the collimator lens L1, as well as the two groups of relay lenses (L3-L4 and L5-L6), were carefully designed so that the Gaussian beam width and the Rayleigh range of the scanned light-sheet were ~9 μm and ~490 μm for 10x configuration. As for 20x configuration, these beam parameters were ~4.5 μm and ~122 μm, respectively. The axial magnification *M*_*axial*_ from the sample plane to IP could be calculated as 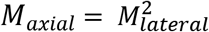 (28). To ensure equal magnification on the image plane in the camera (Andor: Zyla 4.2, 2448×2048 pixels, 6.5 μm pixel pitch), we inserted anamorphic telescope composed of cylindrical lens L7 and L8 between OL3 and L9 (Navitar: MVL75M1, *f* = 75 mm), as described in (25). The anamorphic telescope has optical power only in one dimension such that it can expand the axial dimension by 4x. As the *M*_*axial*_/*M*_*lateral*_ (0.25) in 10x configuration was the reciprocal of the the magnification (4x) of the anamorphic telescope, the magnification difference of *M*_*axial*_ and *M*_*lateral*_ was corrected. Thus the magnifications of the whole optical system along three directions were universially ~2. As for 20x configuration, the later and axial magnification were 4 and 8, respectively.

### 2.2 Optical principle for mesoscopic OPM with low NA objective lens

Fig. 1(b) illustrates the main idea lies behind our design that the angle of the image plane can be increased significantly under Scheimpflug condition to overcome the problem induced by employing a low NA objective lens in mesoscopic scanning OPM.

The relationship of the angle of the illumination plane and IP has been derived in our previous publication (25), thus the angle *θ*_*IP*_ of IP with respect to the optical axis of OA4 is calculated as:

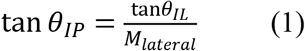

where *θ*_*IL*_ is the angle of the oblique illumination plane with respect to OA1. As the magnification of the two relay lens groups is 0.5, the *M*_*lateral*_ is equal to 0.5 × *f*_*OL*2_/*f*_*OL*1_, where *f*_*OL*1_ and *f*_*OL*2_ is the focal length of OL1 and OL2, respectively. The angle of the illumination plane can be approximated by *θ*_*IL*_ ≈ sin^−1^(*NA*_*OL*1_) ≈ (*NA*_*OL*1_). By substituting *f*_*OL*2_, *M*_*lateral*_, and *θ*_*IL*_ to equation (1), we can reach:

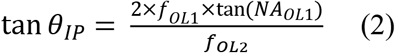

As the value of numerical aperture is small for low NA objective lens, tan(*NA*_*OL*1_) can be approximated by *R*_*OL*1_/*f*_*OL*1_, where *R*_*OL*1_ is the radius of the pupil aperture for OL1. Thus, equation (2) can be reduced to:

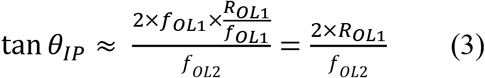

This equation indicates that the angle of IP only depends on the radius of the pupil aperture of OL1 and the focal length of OL2. Therefore, the angle of IP is maintained as long as we choose objective lenses with a similar aperture size. For example, the pupil apertures of low NA Olympus objective lenses (UplanFL4x: *R*_*OL*1_ = 5.8 mm, UplanFL10x: *R*_*OL*1_ = 5.4 mm, and UplanFL20x: *R*_*OL*1_ = 4.5 mm) are close to each other. As a result, the system could be switched into a different configuration with different FOV and resolution by only changing one objective and slightly adjusting the angle and position of the remote imaging system with TS. Specifically for 10× configuration (UplanFL10×/0.3) and 20× configuration (UplanFL20×/0.5) used in this study, *θ*_*IIP*_ is calculated to be ~52° and ~49.1°, respectively.

### 2.3 Imaging protocol

The synchronization of the fast scan mirror GM1, slow scan mirror GM2, and the camera trigger was modified from our previous publication (22,25). Briefly, during every synchronized period, the control system would output one incremental step signal for the slow-scan mirror (GM2), one ramping wave signal for the fast-scan mirror (GM1), and one trigger signal for the camera.

### 2.4 Image processing

As the acquired 2-dimensional images are oblique cross-sectional images, affine transformation was applied to the whole volume data to recover the actual geometry of the sample (22). Geometric transformation functions in Matlab (MathWorks) were used to implement the affine transformation. The color encoded images in depth were generated by ImageJ. For mouse cortex data set, a de-haze method combining the Retinex and wavelet transform algorithm was used to remove the autofluorescence background (29).

### 2.5 Sample preparation protocols

All animal procedures were in accordance with the Institutional Animal Care and Use Committee at Boston Medical Center, Boston University, and conformed to the guidelines on the Use of Animals from the National Institutes of Health (NIH).

Transgenic zebrafish expressing the green fluorescent protein in the entire vasculature Tg(fli1: GFP) were used in the experiment. Zebrafish were cultivated at ~ 28°C following standard procedures. Larvae (≥ 3 days post fertilization, dpf) with a length of 3-4 mm were selected for *in vivo* imaging. The zebrafish larvae were placed in an anesthetic water bath with a regular culture dish during the imaging experiment. The long working distance of low NA objective lens allows imaging through water bath in an upright setup. After the imaging, all of the anesthetized zebrafish were released into freshwater for recovery.

Mouse is perfused transcardially with phosphate-buffered saline (PBS) followed by FITC-albumin (3mg/ml) in gelatin (10%) mixture. The carcass was kept in crushed ice for 10 minutes for the gelatin to solidify. Brain was extracted promptly and immersed in 4% paraformaldehyde (PFA) solution for 6 h and PBS solution for 3 days on a shaker. Tissue is transferred to solutions containing 0.5% alpha-thioglycerol with increasing fructose concentrations for the defined durations (20%: 2-4h, 40%, 60%, 80%:12h, 110%:24h). The brain is then mounted upside down inside a glass bottom Petri dish in the last (110%) solution. It was imaged through the glass bottom.

The transgenic C. elegans strain used in our experiment was QW1217 which pan-neuronally expresses nuclear-localized RFP and cytoplasmic GCaMP6s. Four C. elegans worms were selected 5 days after the hatching stage so that the worms would have a length of ~ 1mm. The worms were placed on the microscope slide and immobilized following a hydrogel encapsulation protocol for imaging.

## 3 Results

### 3.1 Characterization of FOV and resolution

To characterize the resolution and the FOV, phantom measurements were performed. Fluorescent microspheres with a diameter of 3.1 μm were diluted and immobilized in 1% agarose gel. Imaging experiments were carried out with two different objectives with 10x (0.3NA) and 20x (0.5NA) as described in Methods section. The results of the 10x (0.3NA) objective lens are shown in Fig. 2 while the data from 20x can be found in supplementary materials (Fig. S1). Fig. 2(a-c) are the maximum intensity projections (MIP) of the reconstructed data on three orthogonal facets, respectively. The magnifications in X, Y, and Z were calibrated to be 2.02, 2.06, and 2.05, respectively, which agree well with our optical design described above. Given the magnification and the camera’s pitch size (6.5 μm), the FOV along X, Y, and Z direction were measured to be 5.8 mm, 4.9 mm, and 0.7 mm, respectively. Fig. 2(d-f) are zoom-in of the squared area in Fig. 2(a-c).

**Fig. 2.**
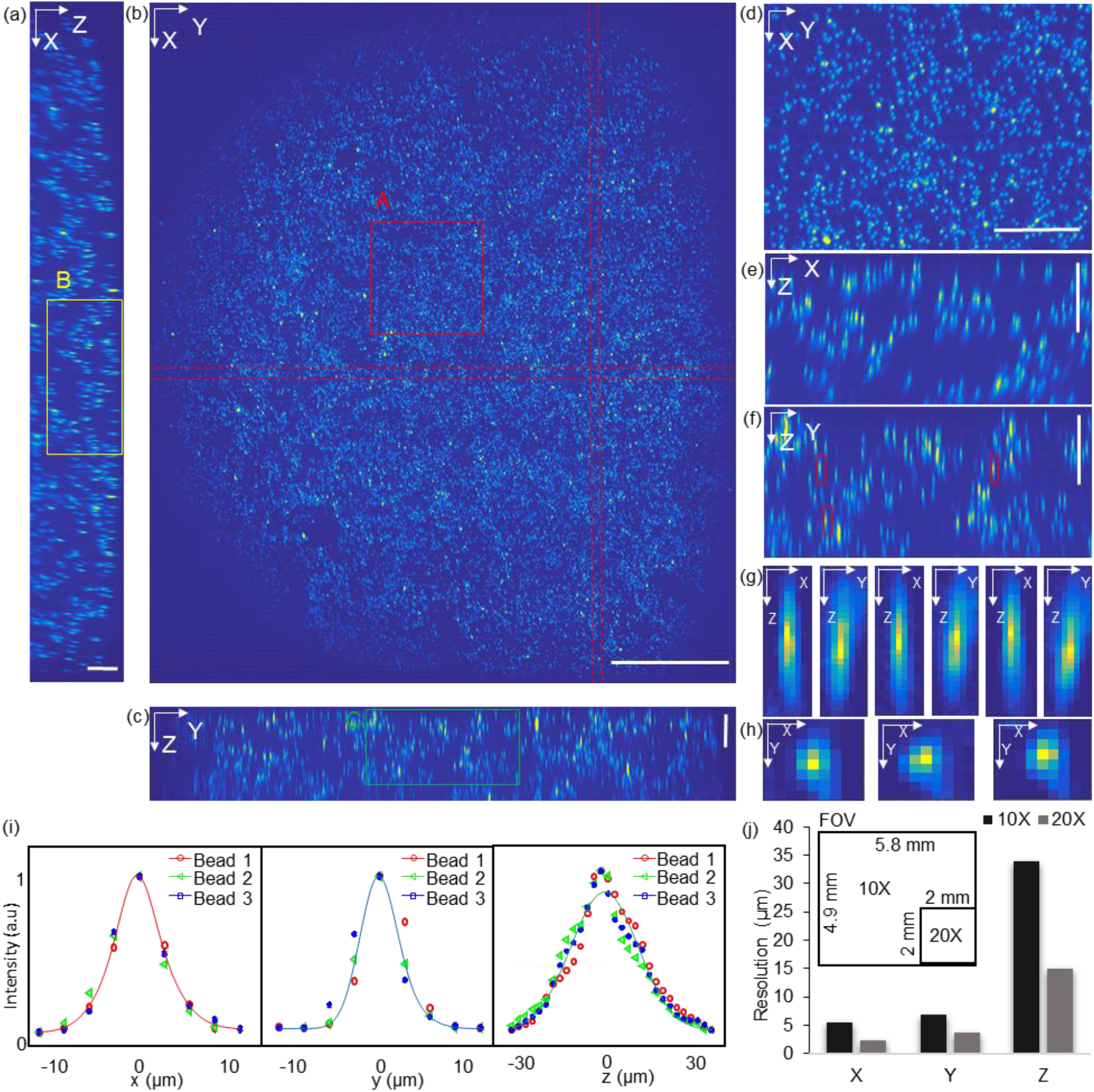
Resolution characterization of mesoscopic OPM. (a-c) Maximum intensity projections of the volume along X, Y, and Z axis. Scale bar in panel (a) and (c) is 0.25 mm. Scale bar in panel (b) is 1 mm; (d-f) The zoom-in view of the squared area A, B, and C in panel (a-c). Both vertical and horizontal bars are 0.25 mm. (g-h) Axial and transversal profiles of three representative beads that are marked in panel (f). (i) Gaussian curve fitting of intensity profile plotting through the center of each bead along x, y, and z direction. (j) FOV and resolution comparison of 10X and 20X configuration.

Three representative beads, which were labeled in Fig. 2(f), were randomly selected to measure the resolutions. The MIPs over each dimension of the three beads are shown in Fig. 2 (g-h). Fig. 2(i) is the intensity distribution through the center of each bead along three dimensions. The intensity data were fitted to the Gaussian curve to quantify the full width at half maximum (FWHM) along each dimension, which was 5.4 μm (X), 6.9 μm (Y), and 34.5 μm (Z), respectively.

Fig. 2(j) is a comparison of both FOV and resolution of 10X and 20X configuration, which show the FOV of 10X configurations is at least 7 times larger than that of 20X configuration while the resolution of the 20X objective is two times better.

### 3.2 Volumetric imaging of *in vivo* zebrafish larvae and *ex vivo* mouse cortex at 10X configuration

Zebrafish is a popular model in biomedical research due to its optical transparency from embryo to larval stages. As the length of the zebrafish larvae is usually longer than 3 mm (≥ 3 dpf), the whole-body volumetric imaging without stitching is difficult by microscopic FOV. Here, transgenic zebrafish expressing green fluorescent protein (≥ 3 dpf) were imaged *in vivo*. The longitudinal body axis of the zebrafish was aligned to be parallel to the light sheet to maximize imaging efficiency. The exposure time of the camera was 14 ms resulting in a frame rate of ~71. The whole imaging process lasted for ~ 3 seconds under 1.2 mW laser power with 220 frames captured by a 3.15 μm slow-scanning step. The size of the acquired volume is 3.2×0.63×0.69 mm^3^ with a pixel dimension 995×218×200. Figure 3(a-c) are MIPs of the whole volume along each dimension. The vascular network can be visulzed in a depth-encoded *en face* MIP in Fig. 3(b). The dorsal longitudinal anastomotic vessels (DLAVs), as well as intersegmental arteries (ISA) and intersegmental veins (ISV) in zebrafish trunk vasculature, is clearly present, each with vessel diameter of ~10 μm (30,31). The space between the paired ISA and ISV is 30 μm to 40 μm (30,31), which can be well resolved in 3D as shown in Fig. 3(b) (lateral) as well as in Fig. 3(c) (axial view / dorsal view). The discrimination of the contra-lateral parachordal vessels (ISA/ISV pair) served as a good demonstration of the axis resolution in our low NA mesoscopic scanning OPM. Figure. 3(a) is a cross-section in the area of the common cardinal vein and the sinus venosus. This feature can also be observed in Fig. 3(d) which are color rendered images in different depth. The structural features presented in each depth is distinct through a total depth of 500 μm.

**Fig. 3.**
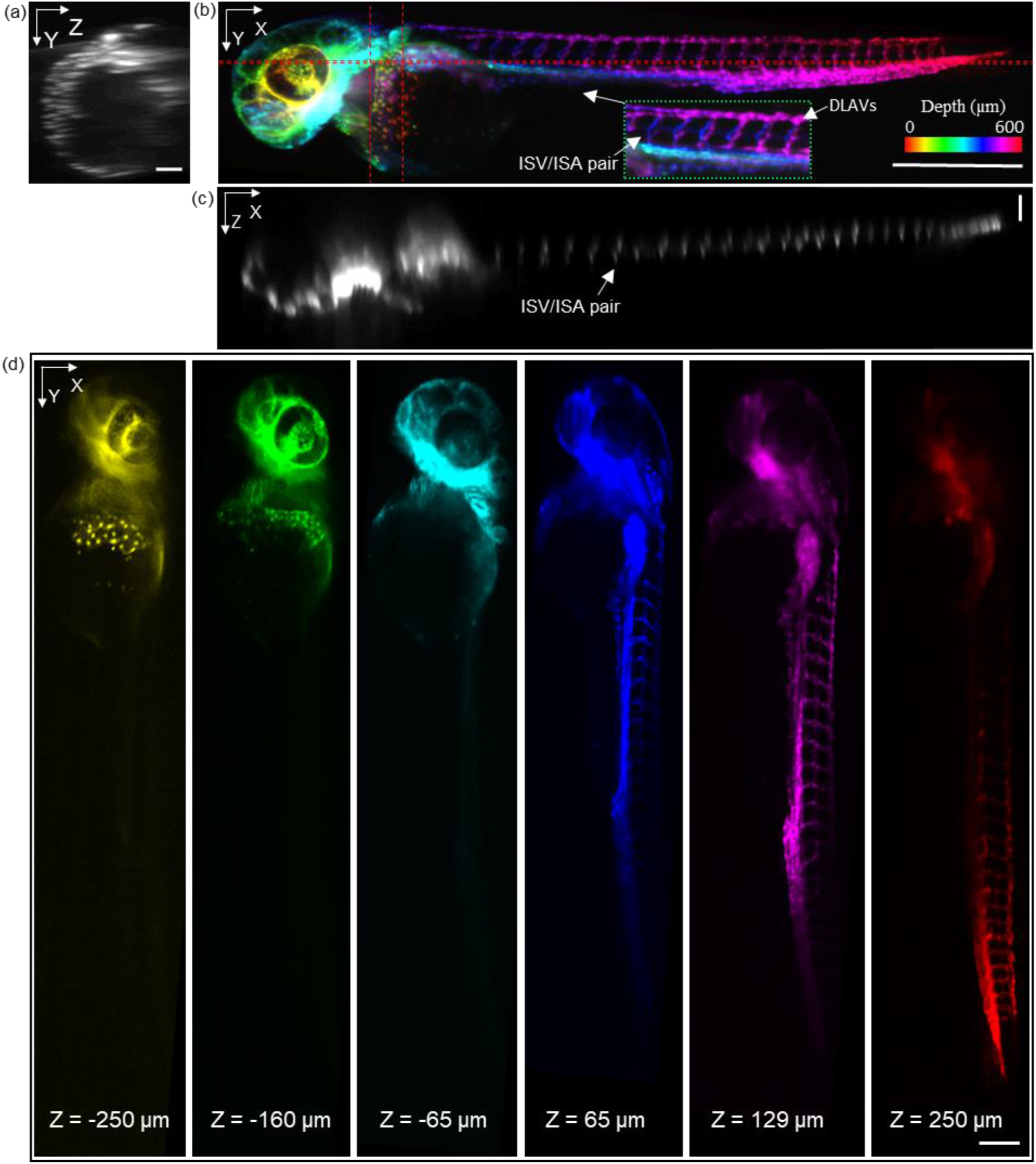
In vivo volumetric imaging of a whole zebrafish larvae. (a, c) Maximum intensity projection (MIP) of the selected layers between the dash lines in panel (b). Both vertical and horizontal scale bars are 100 μm. (b) Depth encoded *en face* view of the whole volume. DLAVs: the dorsal longitudinal anastomotic vessels; ISV: intersegmental veins; ISA: intersegmental arteries; The scale bar is 500 μm. (d) Depth encoded *en face* view in deferent depth. The scale bar is 200 μm.

To demonstrate the extended mesoscopic FOV of the proposed setup, an *ex vivo* mouse cortex with FITC-infused vasculature was imaged with a 10x objective lens. The whole volume was acquired under 0.9 mW laser power with 1550 frames captured by a 3.15 μm slow-scanning step. The resulting volume size is 5.8×4.9×0.7 mm^3^ with a pixel dimension of 1800×1550×220 [Visualization 1]. The depth-encoded *en face* image is shown in Fig. 4(a) in which serval major blood vessels span across the whole FOV as well as their smaller branches. Fig. 4(a) also clearly reveals that the penetrating vessels(PVs) in the mouse cortex, plunging vertically into a deeper layer. Penetrating vessel structure is one of the main vascular topologies in mouse cortex that supports the blood to the neocortex (32). Fig. 4(b) displays the cross-section with color-encoded projection along Y-axis. Penetrating vessels with different colors, which represent the different locations in the Y direction, can be clearly observed. The resolution and image quality is well maintained over the curvature across the FOV (~0.5 mm in depth), thanks to the extended depth of view by the low NA objective lens and the weakly focused excitation. The penetration depth into the highly scattering cortex is estimated to around 0.15-0.25mm based on the PVs.

**Fig. 4.**
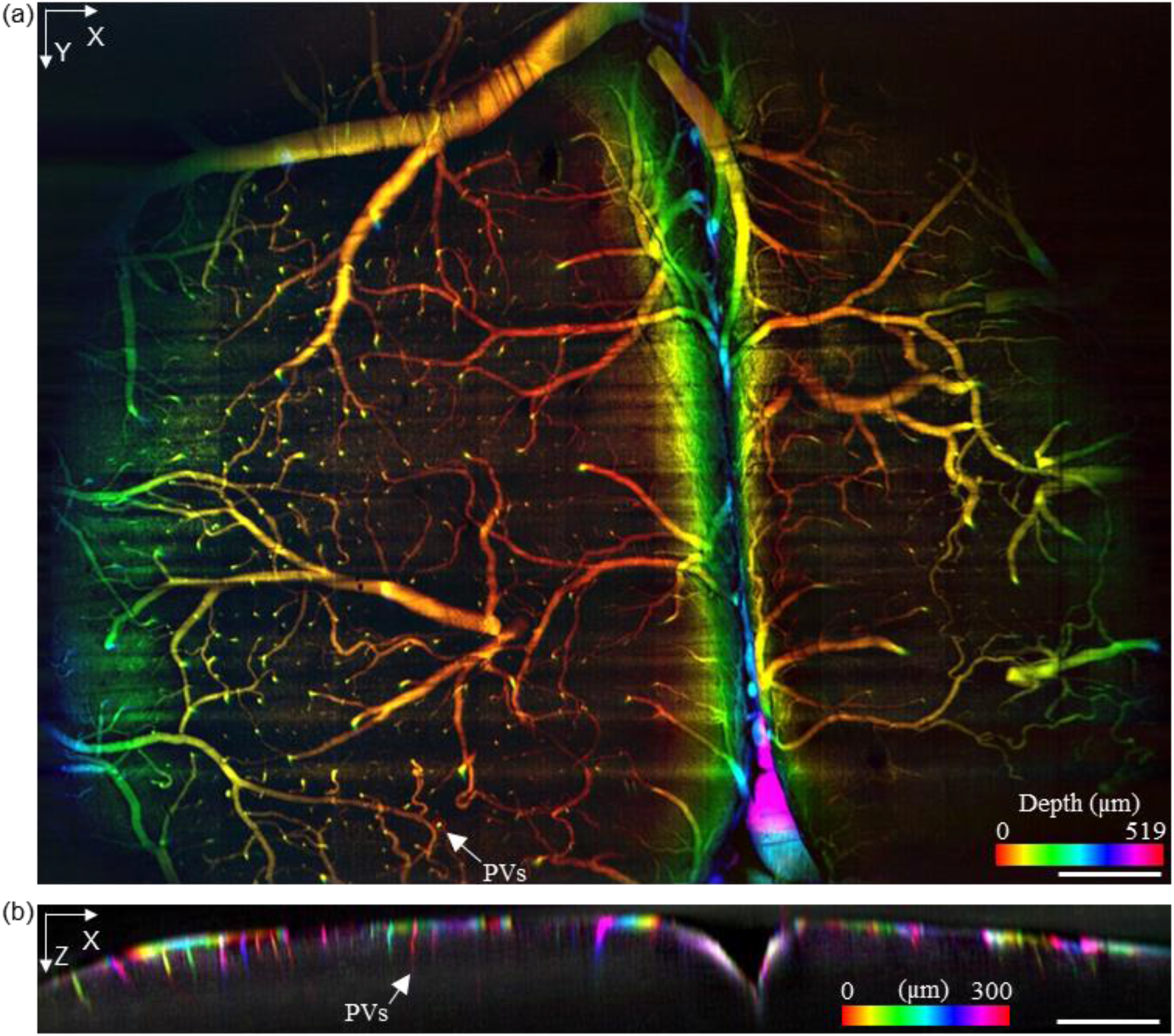
Volumetric imaging of the vasculature in a fixed mouse brain. (a) Depth encoded *en face* view of the mouse brain. (b) X-Z Cross-section with color encoded along Y axis. The scale bar is 500 μm. PVs: penetrating vessels.

### 3.3 Imaging experiment with 20X configuration

By switching to a 20X objective lens and minor refocusing of the remote focusing system, we can flexibly zoom-in a smaller FOV with twice higher 3D resolutions (2.4 μm, 3 μm, 15μm, in *x*-*y*-*z* respectively). Figure 5 demonstrated *in vivo* volumetric imaging of four *C.elegans* expressing GCaMP6s appearing within the same FOV. Fig. 5(a-c) are MIPs along each dimension in which Fig. 5(c) is depth-encoded. Neurons located in various ganglia in the head and tail, along the ventral cord, and inside the body are visible. The excitation laser power was 0.6 mW. 1000 frames were acquired at a slow-scanning step of 1.17 μm. The size of the acquired volume is 1.5×1.2×0.07 mm^3^ with a pixel dimension of 1350×1000×88. Fig. 5(d) is the magnified view in different depths of the selected worm in Fig. 5(c), revealing distinct anatomical features in different depths across the length of the body. Details of another *C. elegan* squared in Fig. 5(c) are shown in Fig. 5(e-h). MIPs of the *C. elegan* are displayed in Fig. 5(e-g), in which the profiles of a worm are visible. Color rendered *en face* views in depth are shown in Fig. 5(h). The individual neurons along the ventral cord and inside the body, and even the thin motor commissures can be observed in different depths.

**Fig. 5.**
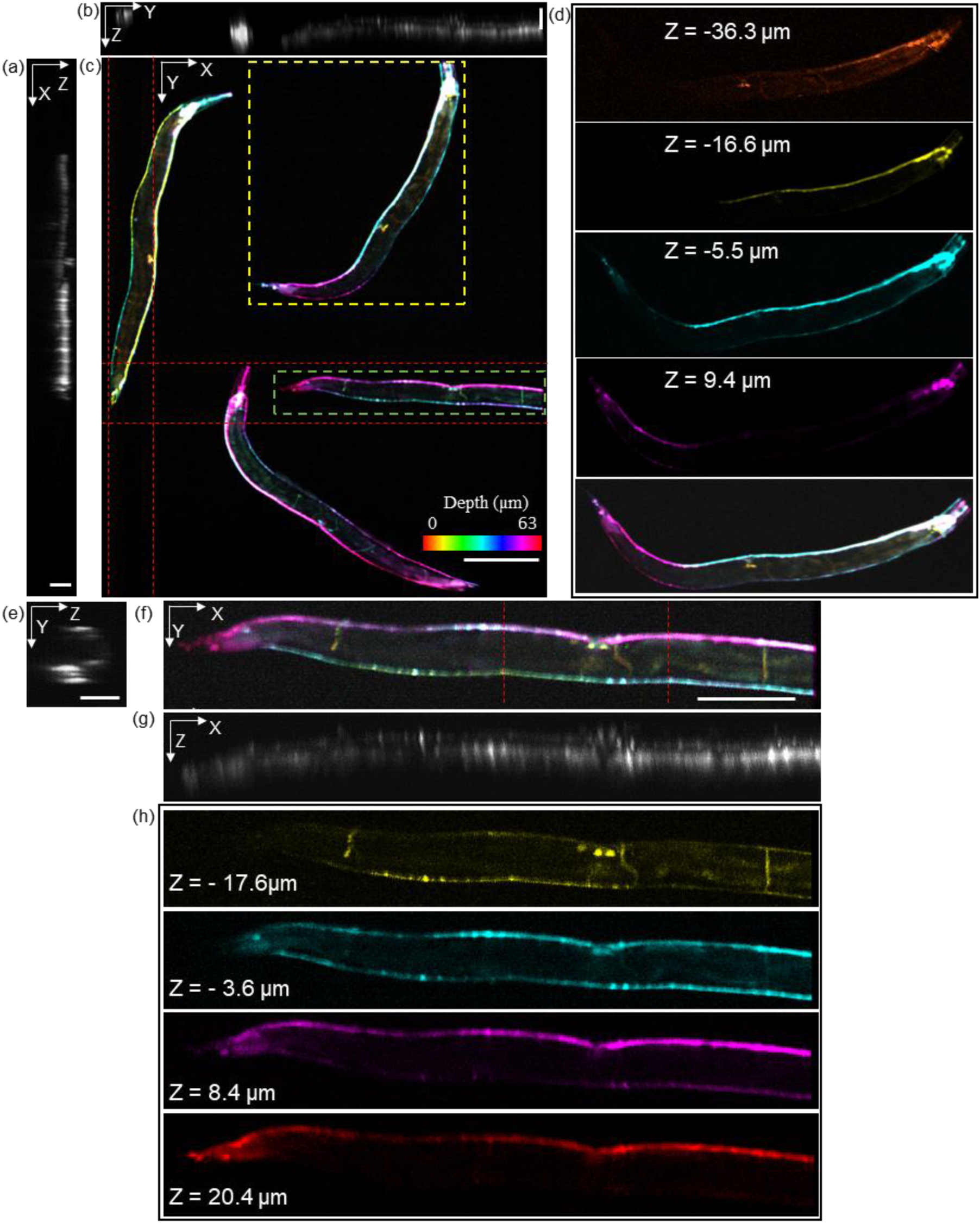
In vivo volumetric imaging of C. elegans worms. (a, b) Maximum intensity projection (MIP) of the layers between the dash lines in panel (c) along Y and X direction. The scale bar is 60μm. (c) Depth encoded *en face* view of the whole volume. The scale bar is 200 μm. (d) Depth encoded *en face* view of the squared worm in panel (c). (e) MIP of the layers between the dash lines in panel (f) along X direction. The scale bar is 30μm. (f) Depth encoded *en face* view of the single worm that squared in panel (c). The scale bar is 100 μm. (g) MIP of the whole worm along Y direction. (h) Depth encoded *en face* view in deferent depths.

## 4 Discussion

We present a scanning OPM design that can achieve mesoscopic volumetric imaging up to ~5×6×0.7 mm^3^ FOV. The proposed method maintains the angle of the intermediate image under Scheimpflug condition by demagnification, and thus allows the use of low NA objective lenses. As compared to the previously reported results (9–22), the achievable FOV (5.8 mm×4.9 mm×0.7 mm) is an order of magnitude higher, the largest FOV in OPM so far to the best of our knowledge. Depth discrimination of vascular structure in zebrafish larvae, mouse cortex, as well as the neurons in *C. elegans*, were demonstrated. Our optical design is versatile that different FOV with varying resolutions can be easily switched by simply changing the objective lenses. Thanks to the single objective lens layout, most common sample formats can be accepted in the proposed method.

As the issue of unparallel light-sheets lying in the scanning geometry of a typical OPM, the tilt-invariant scanning method (19) can be adopted in future improvements. Another major issue of OPM is the light loss in the remote imaging system. Improvements in light collection efficiency can be made by applying water immersion as described in (18). As every pixel represents 3.17 μm in the axial direction of the 10x configuration which is a tenth of the resolution in this dimension, the sampling is redundant in the Z direction in the current setup. By implementing a more suitable sampling scheme, the imaging speed will improve significantly. Due to the highly scattering nature of the mouse brain, we only demonstrated ~0.25mm depth penetration. By switching the Gaussian beam to the Bessel beam or two-photon excitation, the deeper cortex layer could be better revealed. Optical aberrations such as field curvature due to large FOV can be observed in Fig. 2(a), which can be improved and corrected by employing adaptive optics (33).

With the mesoscopic FOV volumetric imaging capability, simple sample mounting protocol, and the versatility of changeable FOVs/resolutions, the proposed method will be ready for the future applications requiring *in vivo* volumetric imaging over a large length scale, such as neural dynamics and vasculature development.

## Supporting information

Visualization 2

## Acknowledgments

This study is supported by National Institute of Health(R01CA224911, R01CA232015, R01NS108464, andR21EY029412) and Bright Focus Foundation (G2017077and M2018132).

## Supplementary Material

**Fig. S1.**
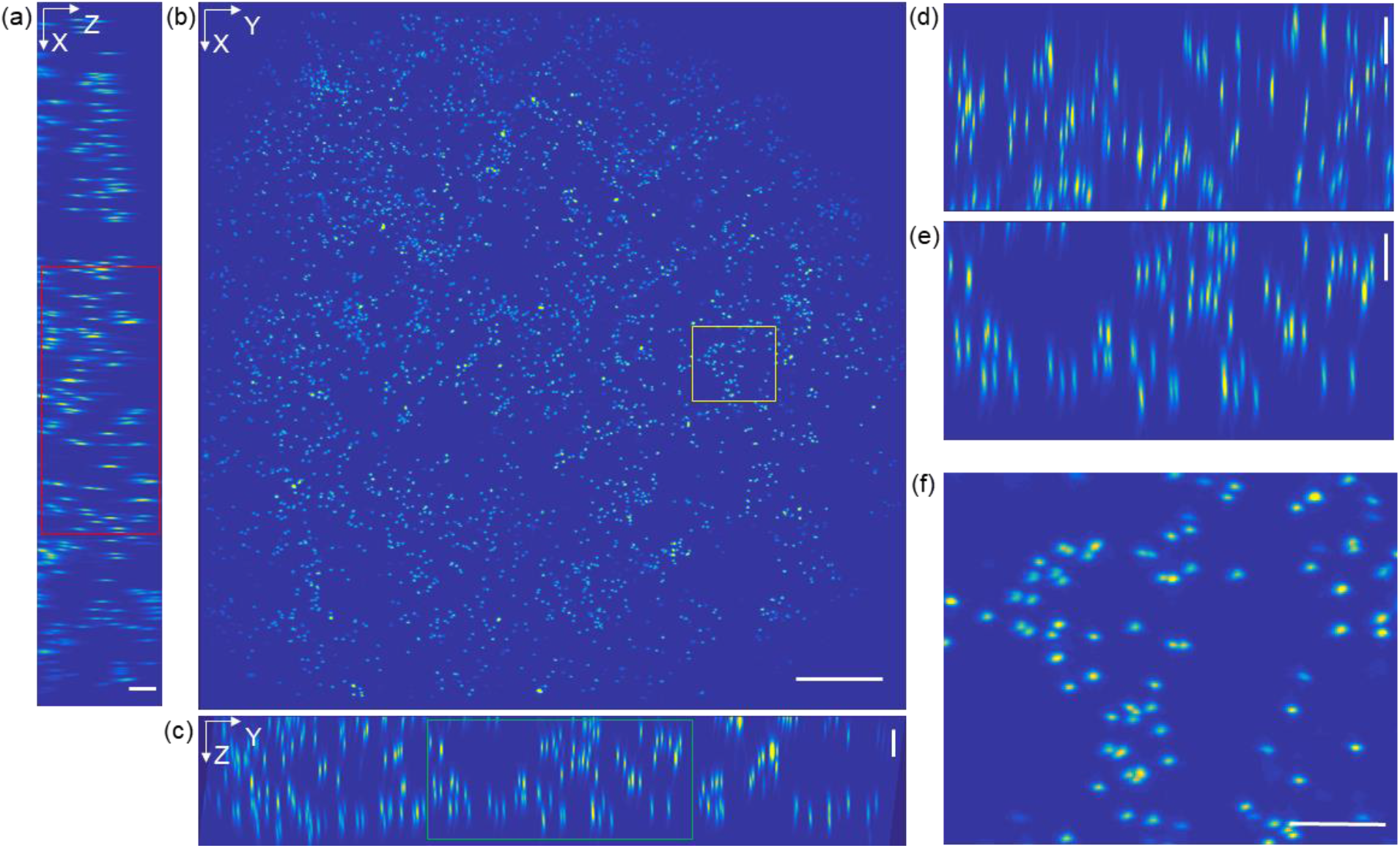
Resolution characterization of 20x configuration. (a-c) Maximum intensity projections of the volume along X, Y, and Z axis. Scale bar in panel (a) and (c) is 50 μm. Scale bar in panel (b) is 250 μm; (d-f) The zoom-in view of the squared area A, B, and C in panel (a-c). Both vertical and horizontal bars are 50 μm.

